# Using neural networks to mine text and predict metabolic traits for thousands of microbes

**DOI:** 10.1101/2020.09.29.319335

**Authors:** Timothy J. Hackmann

## Abstract

Microbes can metabolize more chemical compounds than any other group of organisms. As a result, their metabolism is of interest to investigators across biology. Despite the interest, information on metabolism of specific microbes is hard to access. Information is buried in text of books and journals, and investigators have no easy way to extract it out. Here we investigate if neural networks can extract out this information and predict metabolic traits. For proof of concept, we predicted two traits: whether microbes carry one type of metabolism (fermentation) or produce one metabolite (acetate). We collected written descriptions of 7,021 species of bacteria and archaea from Bergey’s Manual. We read the descriptions and manually identified (labeled) which species were fermentative or produced acetate. We then trained neural networks to predict these labels. In total, we identified 2,364 species as fermentative, and 1,009 species as also producing acetate. Neural networks could predict which species were fermentative with 97.3% accuracy. Accuracy was even higher (98.6%) when predicting species also producing acetate. We used these predictions to draw phylogenetic trees of species with these traits. The resulting trees were close to the actual trees (drawn using labels). Previous counts of fermentative species are 4-fold lower than our own. For acetate-producing species, they are 100-fold lower. This undercounting confirms past difficulty in extracting metabolic traits from text. Our approach with neural networks can extract information efficiently and accurately. It paves the way for putting more metabolic traits into databases, providing easy access of information by investigators.

## Introduction

Microbes are everywhere and can metabolize a huge array of chemical compounds. This makes their metabolism important to nutrient cycling in the environment (1–3). Their metabolism is also important to symbiotic relationships with other organisms (4, 5) and for synthetic biologists in the lab (6, 7). As such, information on microbial metabolism is of value to investigators throughout biology.

Despite the value, information on microbial metabolism is hard to access. Books and journals are filled with this information, but it remains buried in text. Bergey’s Manual (8), for example, reports metabolic traits for thousands of microbes, but in the form of long written descriptions. Looking up information for a few species is feasible, but in the era of big data, investigators often need information on many species.

Information on metabolic traits would more useful if extracted from text and summarized in a database. To date, there is no fast and accurate way of extracting this type of information. One method is to employ teams of curators to read articles and extract information manually (9–11). This method is slow, and information is likely incomplete. Another method is to use machine learning and extract information computationally (12). This method is fast, but accuracy has not been high enough to be adopted by database curators [see ref. (10)].

The field of machine learning has advanced, and it may now have the accuracy needed to extract metabolic information. Neural networks, one form of machine learning, perform well in extracting other kinds of information from scientific literature (13–17). When given medical abstracts, for example, neural networks can recognize and extract out names of diseases (14, 15). Their success with other tasks suggests use in extracting information, such as metabolic traits, from microbiology literature.

One pitfall of neural networks that they must be trained with examples before they can make predictions. These examples, or training data, must be manually labelled by human curators. If large amounts of training data are required, neural networks have little advantage over manual extraction. It is thus critical to determine how much training data neural networks require to predict metabolic traits.

Here we use neural networks to analyze written descriptions of over 7,000 species of microbes and predict their metabolic traits. For proof of concept, we predicted two traits: whether microbes carried out one type of metabolism (fermentation) or produced one metabolite (acetate). Accuracy in predicting these traits was high (>95%). Further, high accuracy was achieved with only modest amounts of training data (labels for 1,000 species). Our approach paves the way to building large databases of metabolic traits, helping investigators working with big data.

## Results

### Collecting text and labels for thousands of microbes

Our general approach to predicting metabolic traits is outlined in Fig. 1. We obtained text (written descriptions of microbial species) from Bergey’s Manual. From this text, we manually labelled metabolic traits. These labels serve as examples to train the network. After training with labels and text, we used the network to predict metabolic traits.

**Fig. 1.**
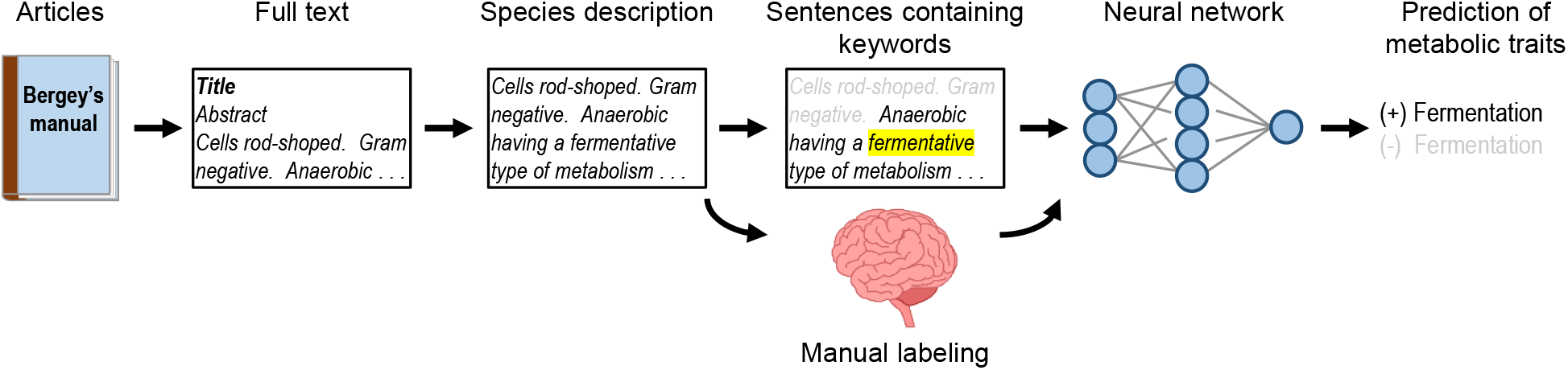
Our approach to predicting metabolic traits with neural networks.

From Bergey’s Manual, we obtained written descriptions for a total of 7,021 species (more exactly, type strains; see list in Table S1). To accomplish this, we downloaded the full text of all genus-level articles (n = 1,503). We extracted out species names, then located relevant sections of text for each species. This extraction was an involved process because names and text for each species were scattered through articles (see Materials and Methods). We assembled the text into coherent species descriptions.

From these descriptions, we manually labelled species as positive or negative for two metabolic traits. The first trait was general: whether microbes carried out one type of metabolism (fermentation). We searched species descriptions for keyword “ferment”. A total of 4,349 descriptions contained the keyword, and we read these descriptions in full. After reading, we labeled species as positive or negative for the trait. Labels (including justifications) are given in Table S1. The second trait was more specific: whether fermentative species produced one metabolite (acetate). We searched for keywords (“ferment” plus “acetate” or “acetic”), read matching descriptions (n = 3,987), then labeled species as positive or negative (see Table S1). Using this approach, we labeled 2,364 species as positive for fermentation, of which 1,009 were also positive for producing acetate. These labels, along with species descriptions, served as training data for the neural network.

### Neural networks accurately predict metabolic traits

After obtaining species descriptions and training data, we trained neural networks and evaluated their performance in predicting metabolic traits. Evaluations were done with data independent from training.

We found neural networks could predict the first metabolic trait (fermentative metabolism) with high accuracy (Fig. 2A). Further, they needed little training data; descriptions for 1,000 species, for example, were enough to achieve 95.3% accuracy. Besides high accuracy, predictions from neural networks achieved high precision and sensitivity (Fig. 2A). Example predictions (from one training with data for 1,000 species) are shown in Table S1.

**Fig. 2.**
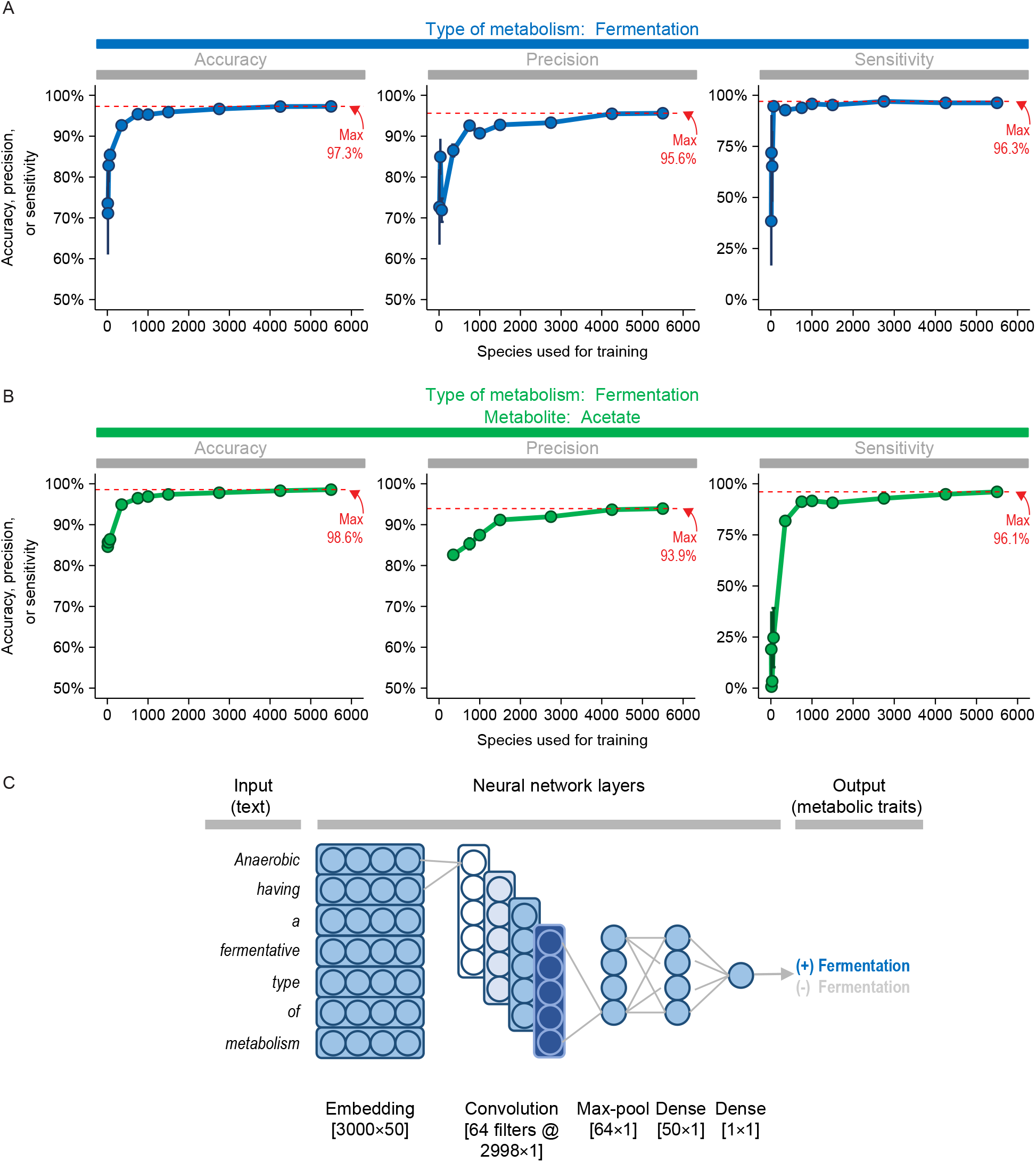
Convolutional neural networks perform well in predicting metabolic traits. (A) Predictions for first trait (fermentative metabolism). (B) Predictions for second trait (acetate production). (C) Architecture of model. Values are means ± SEM of five replicates (independent trainings of the network). Some values for precision are missing because they were undefined (one or more replicates had no false or true positives). For clarity, the number of units depicted in neural network layers is fewer than actual. Units in embedding and hidden dense layers had dropout rate of 0.2.

Neural networks achieved similarly high accuracy when predicting the second trait (acetate production) (Fig. 2B). As before, little training data was needed, and written descriptions of 1,000 species achieved 96.9% accuracy. Precision and sensitivity were also high. Example predictions are shown in Table S1. In sum, neural networks could predict both general and specific traits with modest amounts of training data.

Results above are for the best type of neural network. This type was a convolutional network with architecture shown in shown in Fig. 2C. We tried other types of networks, and a long short-term memory (LSTM) network also performed well (Fig. 3). When little training data was used, its performance equaled or even exceeded that of the convolutional network. However, its performance was overtaken by the convolutional network when using more training data.

**Fig. 3.**
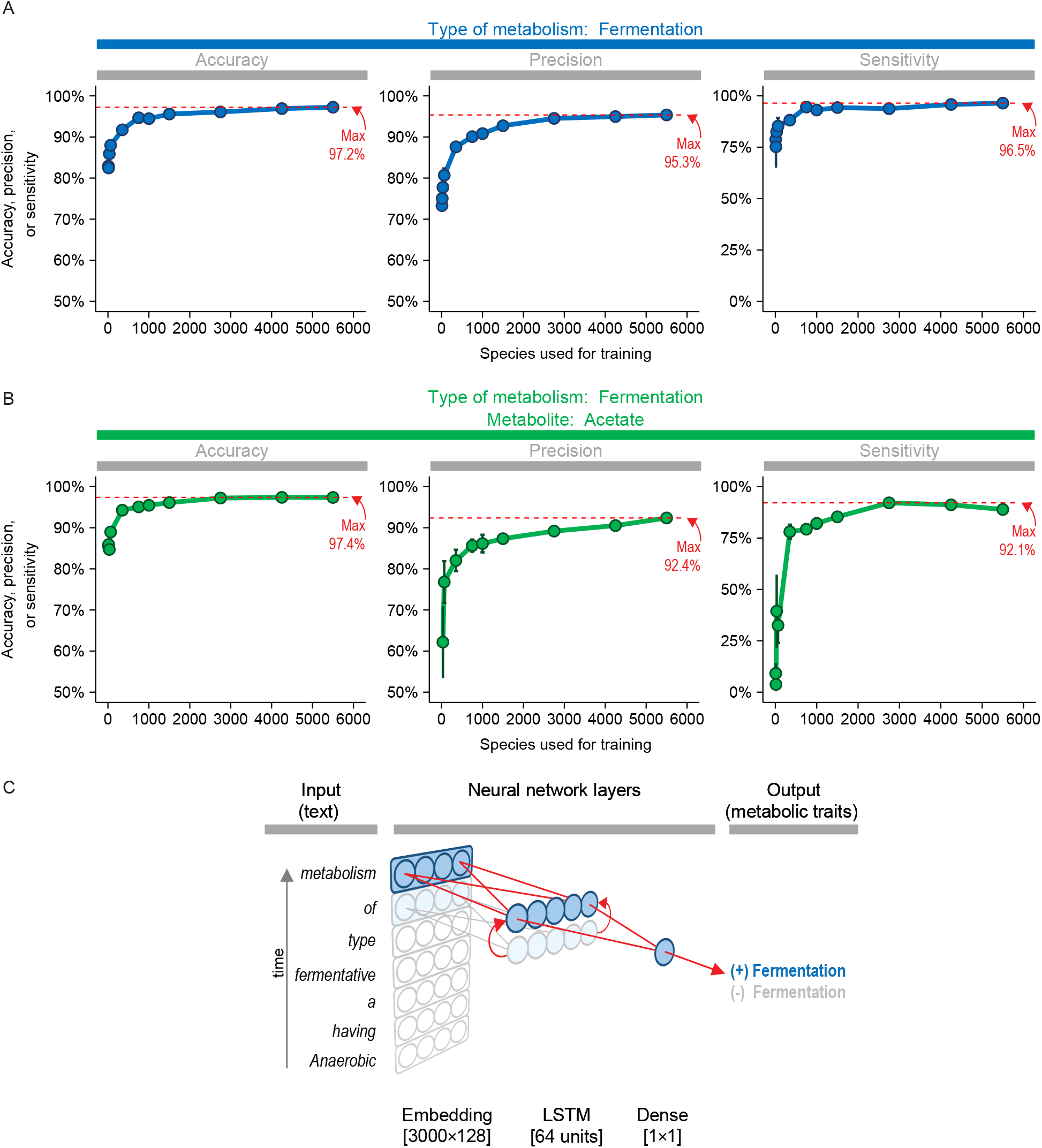
Long short-term memory (LSTM) networks also perform well in predicting traits, though not at the same level as convolutional neural networks. As Fig. 2, except type of network is LSTM. Units in the LSTM layer had a dropout rate of 0.2.

Performance depended not only on the type of network, but also how the text was processed before inputted into the network. The highest performance (shown in Fig. 2) was achieved when the text (species description) was winnowed down to sentences matching key words (e.g., “ferment”). If the full text was used, much more training data was needed (Fig. S1), and performance was never as high. We have thus taken several steps to optimize the network and ensure predictions of metabolic traits are as high as possible.

### Predictions from neural networks yield accurate phylogenetic trees

We evaluated neural networks further by drawing phylogenetic trees of their predictions. First, we made a phylogenetic tree of all species in Bergey’s Manual (Fig. 4A). Next, we highlighted species predicted to have the first trait (fermentative metabolism) (Fig. 4B). We found the predicted species were similar to those manually labeled as having it. We found similar agreement between predictions and labels for the second trait (acetate production) (Fig. 4C). For both traits, we used training data for only 1,000 species. In sum, predictions from neural networks were not just accurate in a statistical sense. They produced phylogenetic trees that were close to the actual ones, showing they are accurate biologically.

**Fig. 4.**
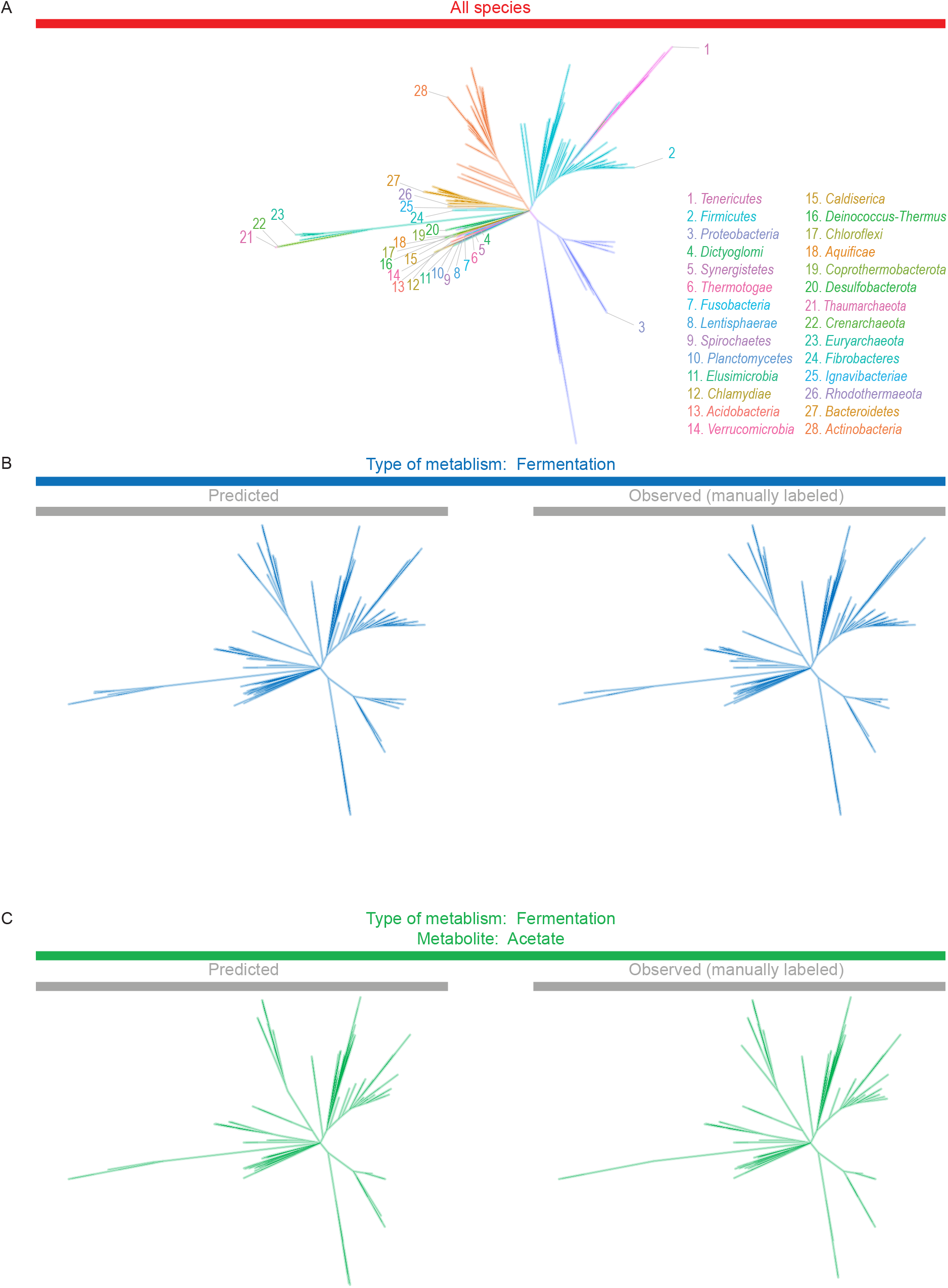
Predictions of neural networks lead to accurate phylogenetic trees. (A) All species in Bergey’s Manual with available sequences. (B) Species with first trait (fermentative metabolism). (C) Species with second trait (acetate production). When traits were predicted, they were done so by convolutional neural network and training data for 1,000 species. Trees were constructed with concatenated ribosomal protein sequences as described in Materials and Methods.

### Databases reporting metabolic traits are incomplete

Some information on metabolic traits can already be found in databases, but it is not clear how complete it is. Because our work identified metabolic traits for a number of species, it can help assess how complete are these databases.

As mentioned, our work identified 2,364 species that carried out fermentation. By comparison, the best database identified 644 species, or 27% of our number (Fig. 5). For species that also produce acetate, the best database identified 1.2% of our number.

**Fig. 5.**
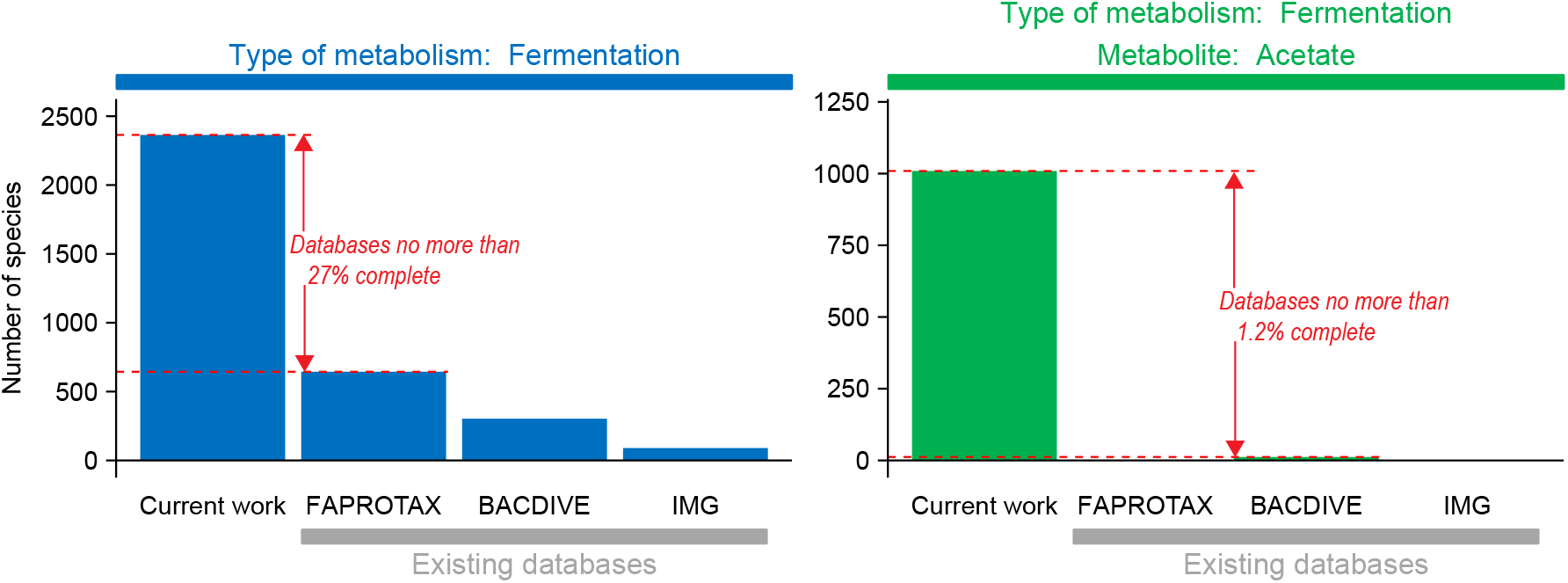
In comparison to the current work, existing databases reporting metabolic traits are incomplete. Species reported to have a metabolic trait were counted as described in Materials and Methods.

Our own numbers of species are incomplete, and thus the situation is even worse than it first appears. We obtained descriptions for 7,021 species, yet the total number of species validly published in the literature is 20,038 [see ref. (18)]. In total, our results suggest that databases reporting metabolic traits are far less than 1/4 complete.

## Discussion

Microbial metabolism cuts across many fields of biology, yet information on metabolic traits is still hard to access. The information is locked away in text of books and articles. Several attempts have been made to extract this information and make it available in databases (9–11, 19, 20). However, the information collected so far appears far less than 1/4 complete (see Fig. 5). Most attempts to extract information have done so manually, using teams of curators (9–11). To provide more complete information, a faster method is needed.

We propose neural networks as a fast (and accurate) method to extract information and predict metabolic traits. We provide proof of concept by predicting two metabolic traits for thousands of microbes and with >95% accuracy. This level of performance was high enough to create an accurate phylogenetic tree of these species, and it should be useful for other applications. The performance of our networks represents an improvement over using other types of machine learning to predict metabolic traits of microbes. Mao et al. (38), for example, predicted traits with a support-vector machine. This approach gave 59% precision and 66% sensitivity when predicting metabolites produced during fermentation. With neural networks, we achieved 93.9% precision and 96.1% sensitivity for a similar prediction (see Fig. 2).

Despite the promise of our approach with neural networks, there are still areas that need to be explored. We need to explore, first, sources of species descriptions other than Bergey’s Manual. Though Bergey’s Manual gave us descriptions for over 7,000 species, this represents only ~1/3 of all species validly published in the literature (18). We need to explore, second, well our methods work with rare traits. Both metabolic traits we investigated were relatively common (found in over 1,000 species).

Once these uncertainties are resolved, neural networks can be deployed at an even larger scale to predict metabolic traits of microbes. They would enable building of databases of metabolic traits larger than previously imagined. These databases, in turn, will be key to opening up the study of microbial metabolism and bringing it fully into the era of big data.

## Materials and Methods

### Preparation of text

To obtain written descriptions of species, articles from Bergey’s Manual were downloaded and read into R. Names of species were extracted from the full text, then appropriate sections of the full text were assembled into the description. Full code for preparing text is available at https://github.com/thackmann/MicroMetabolism.

Articles in Bergey’s Manual were downloaded as html files. This was done using article urls in Browse A-Z page in Bergey’s Manual and the download.file() function in R. Only genus-level articles (containing “gbm” in their url) were retained.

The html files were read into R. The full text of each article was then obtained using html_nodes() function and css selectors.

Names of each species were extracted from the full text. For a given article, the genus name was extracted using css selectors. Names of species were then found under the List of Species of the Genus section using the genus name and regular expressions. We reviewed the list of names manually, identified errors, and refined regular expressions (using different expressions to accommodate varying format of articles). Our list also included names of subspecies, biovars, pathovars, and genomospecies, which we treated as equal to species. A similar approach was used to extract other taxonomic ranks and strain IDs.

The full text was parsed to give a written description of each species. The full text typically consisted of 1) Abstract, 2) Further Descriptive Information and other sections about the genus, 3) List of Species of the Genus, and 4) References. These sections were identified using regular expressions. For a given species, we combined text from sections (2) and (3). For (3), we selected only text belonging to the given species, and we excluded text for other species within the genus. This text was selected by using regular expressions for the species name.

### Labeling of metabolic traits

We labeled species as positive or negative for two metabolic traits. Using R and regular expressions, we searched the species descriptions for keywords. For the first trait (fermentative metabolism), the keyword was “ferment”. For the second trait (acetate production), the keywords were “ferment” plus “acetate” or “acetic”. The regular expression allowed matches not just to the keyword itself, but to any word containing it. For the keyword “ferment”, the words “ferment”, “fermenter”, and “non-fermentative” would all match. When there was a match to the keyword, we read species descriptions in full before labeling the species as positive or negative for the trait. If there was no match, the species was labeled as negative.

### Construction of neural networks

Neural networks were built in R using functions in the Keras library. Full code for constructing neural networks is available at https://github.com/thackmann/MicroMetabolism.

Written description of each species were prepared for input into the network. Sentences matching the keywords were kept, and others were discarded. For the first trait (fermentative metabolism), the keyword was “ferment”. For the second trait (acetate production), the keywords were “ferment” plus “acetate” or “acetic”. At least one sentence had to match “ferment” for any to be kept. Some sentences were duplicated, and these were discarded. The remaining sentences were joined together and truncated at 25,000 characters. Afterwards, the text was tokenized using the text_tokenizer(), fit_text_tokenizer(), and texts_to_sequences() functions with num_words of 3,000. The tokenized text was then inputted into the network as a list with one element per species.

Labels of metabolic traits were inputted as a vector with one element per species. The elements were 1 (trait positive) or 0 (trait negative).

The networks had architecture as shown in Fig. 2 and 3. They were solved with the loss function binary_crossentropy and adam optimizer. The networks were trained with batch size of 32 for 10 epochs. For small amounts of training data, more epochs (up to 40) were needed to minimize the loss function. The amount of training data was as specified in Fig. 2 and 3.

Predictions were evaluated using accuracy, precision, and sensitivity. Accuracy was calculated as (TP+TN)/(TP+TN+FP+FN), where TP = true positive, TN = true negative, FP = false positive, and FN = false negative. Precision was calculated as TP/(TP+FP). Sensitivity was calculated as (TP)/(TP + FN).

### Construction of phylogenetic trees

We constructed a phylogenetic tree of genomes belonging to species from Bergey’s Manual. The construction followed the general approach of ref. (21, 22) and used sequences of 14 ribosomal proteins.

First, we used the strain IDs of each species to find genome sequences. Specifically, we used the strain ID to find a GOLD organism ID (23), GOLD project ID (23), and the IMG/M genome ID (genome sequence) (19) (see Table S1). Though we could have searched IMG/M directly with the strain ID, this approach was slow. Some strain IDs were generic (e.g., numbers like “238”) and could match multiple GOLD organism IDs. To make matches more specific, we required the species or genus name to match, also. We identified genome IDs for a total of 2,925 species.

Next, we downloaded amino acid sequences of the ribosomal proteins from IMG/M (19). We did this using KO IDs for the respective genes (Table S2) along with IMG/M genome IDs. We discarded sequences that were short (<75% of the average length for a given ribosomal protein).

We aligned sequences with Clustal Omega in R (24–26) and then concatenated them. We discarded columns in the alignment with a large number of gaps (95% or more).

We used aligned and concatenated sequences to create a phylogenetic tree. The tree was calculated using maximum likelihood with RAxML (27) on the CIPRES web server (28). The parameters are listed in Table S3.

Final analysis and visualization were done in R. The consensus tree and branch lengths were calculated using phytools (29). The tree was visualized using ggtree (30). A total of 2,501 species had genomes with protein sequences that could be included in the final tree.

In the full tree, we highlighted branches belonging to species predicted or labeled to have a metabolic trait. These predictions were made using the convolutional neural network in Fig. 2C and training data for 1,000 species. Species part of training data were not highlighted, even if they had the trait.

### Completeness of databases reporting metabolic traits

We investigated the completeness of information in three databases: FAPROTAX (11), BacDive (9), and IMG (19). We did not investigate the IJSEM database (10) because its information has been subsumed by BacDive (9). We also did not investigate the MACADAM database (20) because its information is in FAPROTAX (11) and IJSEM (10) databases.

For the three databases, we counted the number of microbial species they report as having a fermentative metabolism. For FAPROTAX (v. 1.2.3) (11), we counted entries under the functional group “fermentation”. Only entries containing both genus and species names were counted. For BacDive (9), we used Advanced search > Morphology and physiology > Metabolite (utilization). We set Kind of Utilization to “fermentation” and Utilization activity to “+”. For IMG/M (19), genomes with information on metabolism were displayed using Genome Search > Advanced Search Builder > Metabolism. We searched the output for the keyword “ferment” and then read the description in full.

We also counted the number of species the databases reported as producing acetate. For FAPROTAX, we counted no species because no functional group indicated both fermentative metabolism and acetate production. For BacDive, we entered the same settings as for the first trait (fermentative metabolism). Additionally, we set Metabolite (production) to “acetate” and Production to “yes”. For IMG/M (19), we manually searched the output for the keywords “acetate” and “acetic”, then then read the description in full.

### Data availability

Code for preparing text and constructing neural networks is available at https://github.com/thackmann/MicroMetabolism.

## Acknowledgements

This work was supported by a Hatch Project [accession no. 1019985] from the United States Department of Agriculture National Institute of Food and Agriculture.

## Supplemental material legends

*Supplemental tables are not available in the pre-print version of the manuscript.*

**Table S1.** Metabolic traits and other information on species from Bergey’s Manual.

**Table S2.** Ribosomal proteins and database IDs searched.

**Table S3.** Parameters for calculating the phylogenetic tree in RAxML

**Fig. S1.** Performance of neural networks when inputting full text. As Fig. 2, except the full text, not just sentences containing keywords, was inputted. Before tokenization, sentences were truncated to 200,000 instead of 25,000 characters. During tokenization, num_words was set to 5,000 instead of 3,000.

