## Supplementary material for "Using neural networks to mine text and predict metabolic traits for thousands of microbes": Fig. S1

**Supplemental material figure legends**

**Fig. S1.** Performance of neural networks when inputting full text. As Fig. 2, except the full text, not just sentences containing keywords, was inputted. Before tokenization, sentences were truncated to 200,000 instead of 25,000 characters. During tokenization, num_words was set to 5,000 instead of 3,000.

Figure S1

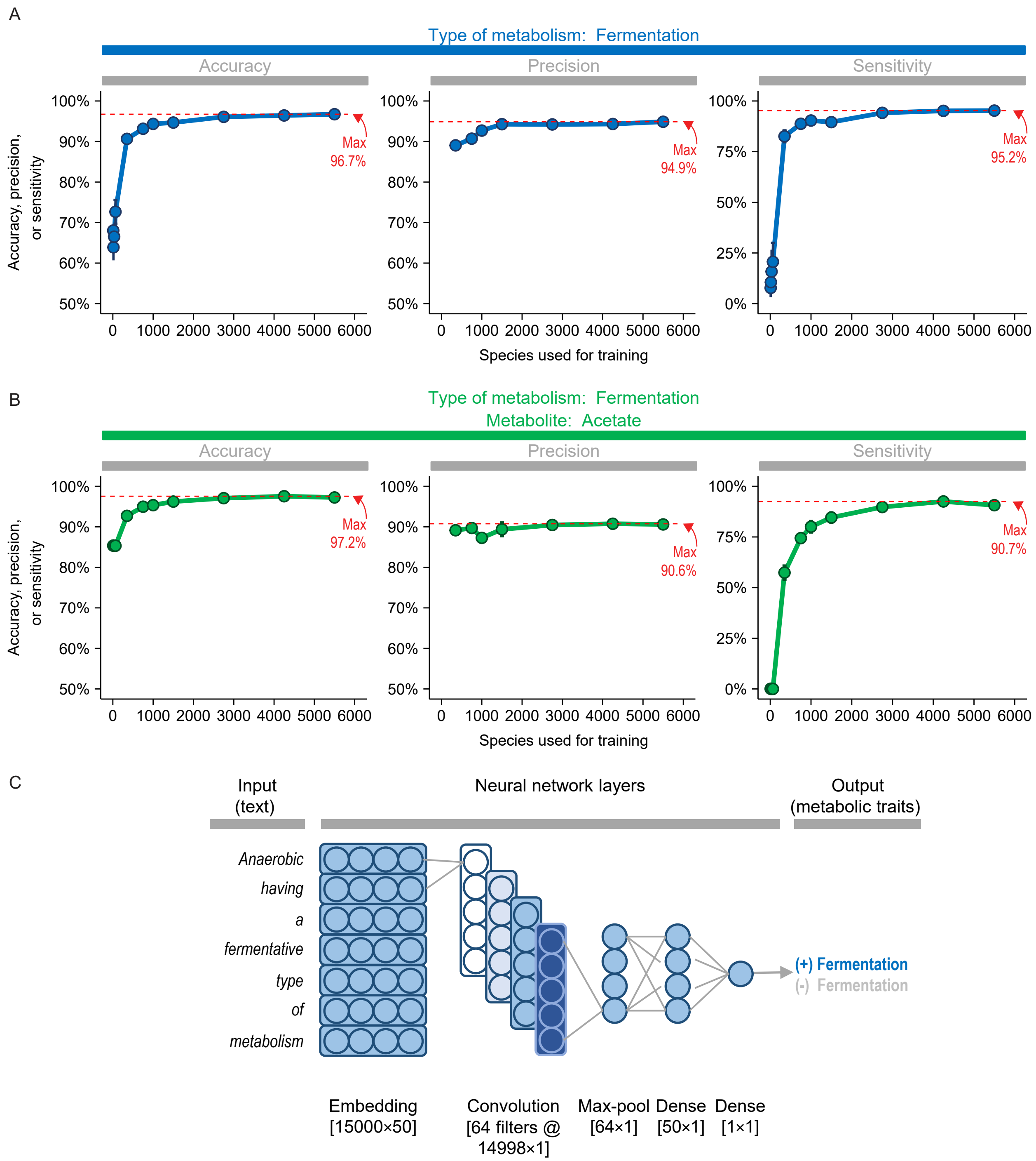
A
Type of metabolism: Fermentation
Accuracy
Precision
Sensitivity
Accuracy, precision, or sensitivity
Max 96.7%
Max 94.9%
Max 95.2%
Species used for training
B
Type of metabolism: Fermentation
Metabolite: Acetate
Accuracy
Precision
Sensitivity
Accuracy, precision, or sensitivity
Max 97.2%
Max 90.6%
Max 90.7%
Species used for training
C
Input (text)
Neural network layers
Output (metabolic traits)
Anaerobic
having
a
fermentative
type
of
metabolism
(+) Fermentation
(-) Fermentation
Embedding [15000×50]
Convolution [64 filters @ 14998×1]
Max-pool [64×1]
Dense [50×1]
Dense [1×1]
